# Environmental enrichment delays the development of stereotypic behavior and reduces variability in behavioral experiments using California mice (*Peromyscus californicus*)

**DOI:** 10.1101/2021.01.05.425454

**Authors:** Vanessa Minie, Stephanie Ramos-Maciel, Emily Wright, Radmila Petric, Brian Trainor, Natalia Duque-Wilckens

**Affiliations:** Department of Psychology, University of California Davis, CA 95616; Department of Physiology, Michigan State University, East Lansing MI 48824; Department of Large Animal Clinical Sciences, Michigan State University, East Lansing, MI 48824; Department of Biology, University of North Carolina at Greensboro, Greensboro, NC, United States

## Abstract

Domesticated mice and rats have shown to be powerful model systems for biomedical research, but there are cases in which the biology of species is a poor match for the hypotheses under study. The California mouse (*Peromyscus californicus*) has unique physiological and behavioral traits and has emerged as a powerful model for studying sex differences in the biology of psychiatric disease, which is particularly relevant considering the new NIH guidelines that require the inclusion of sex as a biological variable. Despite its growing role in preclinical research, there is a lack of studies assessing species-specific housing needs, which presents a challenge for research facilities seeking to ensure good welfare and obtaining high-quality experimental data. Indeed, captive California mice present a high prevalence of stereotypic backflipping behavior, a common consequence of suboptimal housing and a potential source of experimental outcome variability. Using three different cage systems, the present studies show that increasing housing space as well as social and environmental complexity can delay the development of stereotypic behavior in male and female California mice. Critically, this reduction in stereotypy is accompanied by increased effect sizes of stress in an established model for social anxiety. These results suggest that increased cage size and enrichment could enhance welfare in California mice while simultaneously increasing the quality of behavioral experiments.

## Introduction

The California mouse (*Peromyscus californicus*) possesses unique characteristics that make it a powerful model species for testing hypotheses that are difficult or impossible to test with standard rodent lines. First, California mice are monogamous, biparental, and both males and females show aggression towards intruders of both sexes (Ribble and Salvioni, 1990; Rieger et al., 2019). Male parental behavior and female aggression are present in humans but are very rare in common laboratory rats and mice (Kleiman, 1977; Kleiman and Malcolm, 1981), making the California mouse a great model to study the sex-specific mechanisms underlying social behaviors relevant to human behavior. This is particularly important considering that the new NIH guidelines require consideration of sex as a biological variable in research. Second, California mice are prone to developing insulin resistance and hyperlipidemia independent of obesity, which makes this species an ideal model for the study of the early stages of metabolic syndrome (Krugner-Higby et al., 2011, 2006, 2000), one of the major health hazards of the modern world (Saklayen, 2018). Together, this suggests that the use of the California mice as a preclinical model will increase in the coming years, as it has been for the past decade (source: Web of Science).

While the unique physiology and behavior of California mice provide great opportunities for research, they can also present unique challenges for husbandry. Although housing needs can greatly vary among rodent species (Baumans, 2005), there are no species-specific guidelines for California mice (National Research Council (US) Committee for the Update of the Guide for the Care and Use of Laboratory Animals, 2011), which can result in suboptimal housing conditions for this species. Indeed, California mice colonies can present a high prevalence of stereotypic back-flipping (Greenberg et al., 2014), a common consequence of suboptimal housing (Garner, 2005; Gross et al., 2012), and a potential indicator of poor welfare (Mason and Latham, 2004). Importantly, while most individuals in California mouse colonies display this stereotypic behavior at some point, there is great inter-individual variability in the frequency at which this behavior is performed. Stereotypic back-flipping has been reported in multiple species (Mason and Latham, 2004), and it is a problem because stereotypic behavior can be accompanied by altered physiology (McBride and Parker, 2015), cognition (Garner and Mason, 2002), and affective state (Novak et al., 2016). increasing the variability of experimental outcomes (Garner, 2005). We were therefore highly motivated to find housing alternatives that could minimize stereotypic behaviors. With this aim, here we tested three different housing conditions on backflipping behavior, the social interaction test, and welfare indicators in male and female California mice. The housing conditions were a) standard, b) larger cages and increased social complexity (large), and c) larger cages, increased social complexity, and environmental enrichment (large+EE). We designed the enrichment program based on observations of California mice in the wild, who build complex nests in a variety of contexts including tree cavities, leaf litters, and creek banks, and spend a significant amount of the day climbing (Dalquest, 1974; Gubernick and Alberts, 1987). Based on findings in other species (Bayne, 2018; Bayne and Würbel, 2014; Bechard et al., 2016; Shyne, 2006), we hypothesized that large+EE would be the most effective at reducing stereotypic behavior and that this would in turn help reduce variability in the social interaction test.

## Materials and Methods

### Animals and housing

All mice were bred in our colony (University of California, Davis, Department of Psychology). At weaning (post-natal day 30, P30) animals were ear punched for identification and assigned to one of three conditions (fig.1A,B): 1. standard (15cm ×25cm ×12cm cage, 2 same-sex individuals), 2. Large (10.5in ×19in ×6in cage, 4 same-sex individuals), or 3. large+EE (4 same-sex individuals). All cages were made of clear polypropylene and all treatment groups were provided with Sani-chip bedding, cotton nestlets, and enviro-dri (Newco Distributors). The enrichment group was additionally provided with a crawl ball™ (Bio-Serv®), a stainless-steel loft with holes (Otto environmental®), and a 4×6inch stainless-steel tube (Otto environmental®). For all animals, water and food (Harlan Teklad 2016; Madison, WI) were provided ad libitum. All treatment groups were housed in the same room under a 16L:8D light:dark cycle. The room temperature was kept at (20-23°C). All behavior experiments were carried out during the dark cycle (14:00-17:00), the time at which California mice are most active. Experimenters used red light (3 lux) headlamps to help with experimenter vision but minimize potential effects of light exposure on the mice circadian rhythm (Hattar et al., 2003; Provencio and Foster, 1995; Yoshimura and Ebihara, 1996).

**Fig.1.**
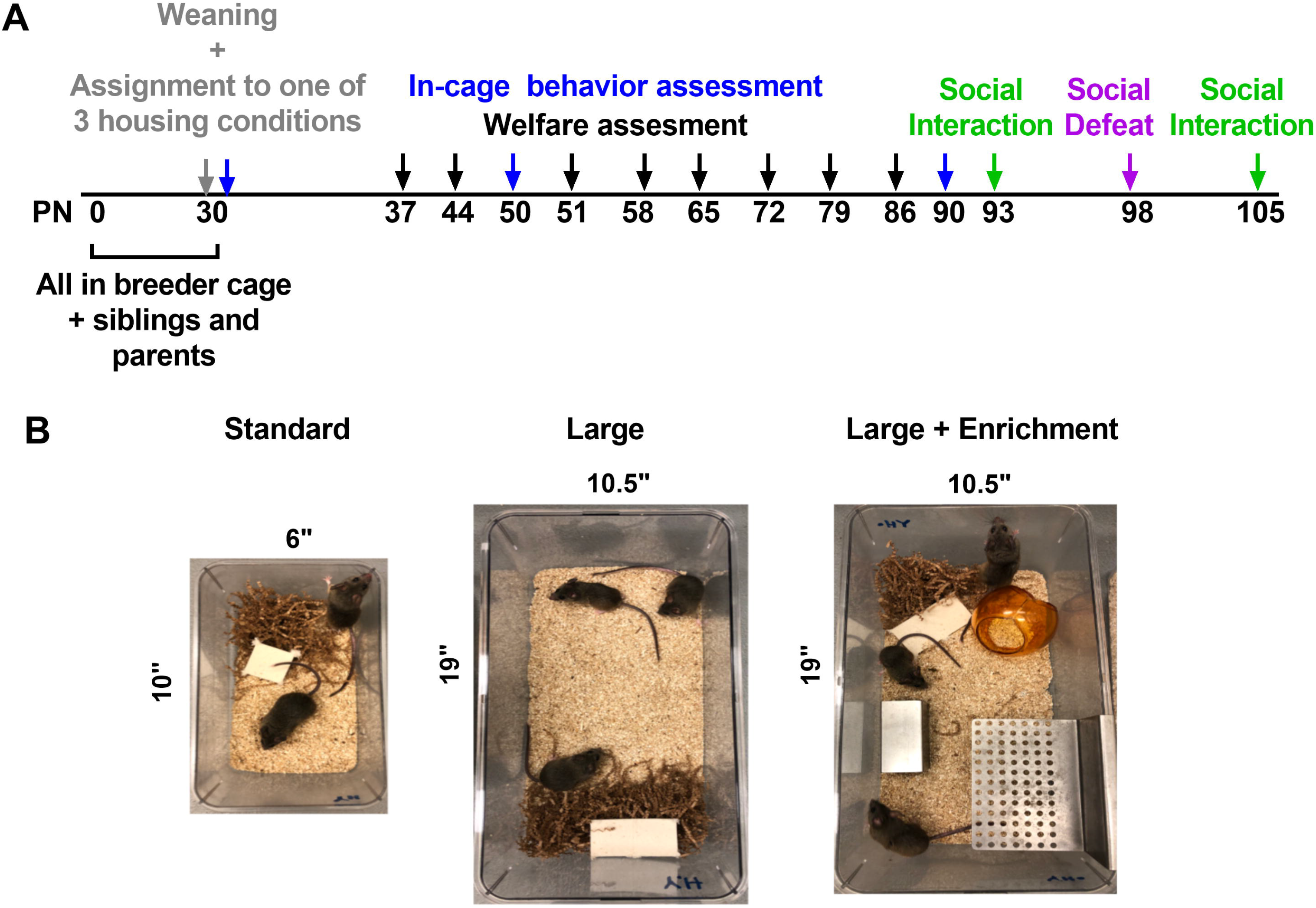
General experimental timeline (A). Pictures showing three experimental housing conditions: standard, large, and large + Enrichment (EE) (B).

### In-cage behavior

Individual in-cage behavior was assessed using one 5 minute observation at 3 developmental stages: P30, P50, and P90 (fig. 2A). Every 15 seconds, the presence/absence of back-flipping was recorded. No other stereotypic-like behaviors were observed. In the large+EE group, the use of enrichment was also recorded.

**Fig. 2.**
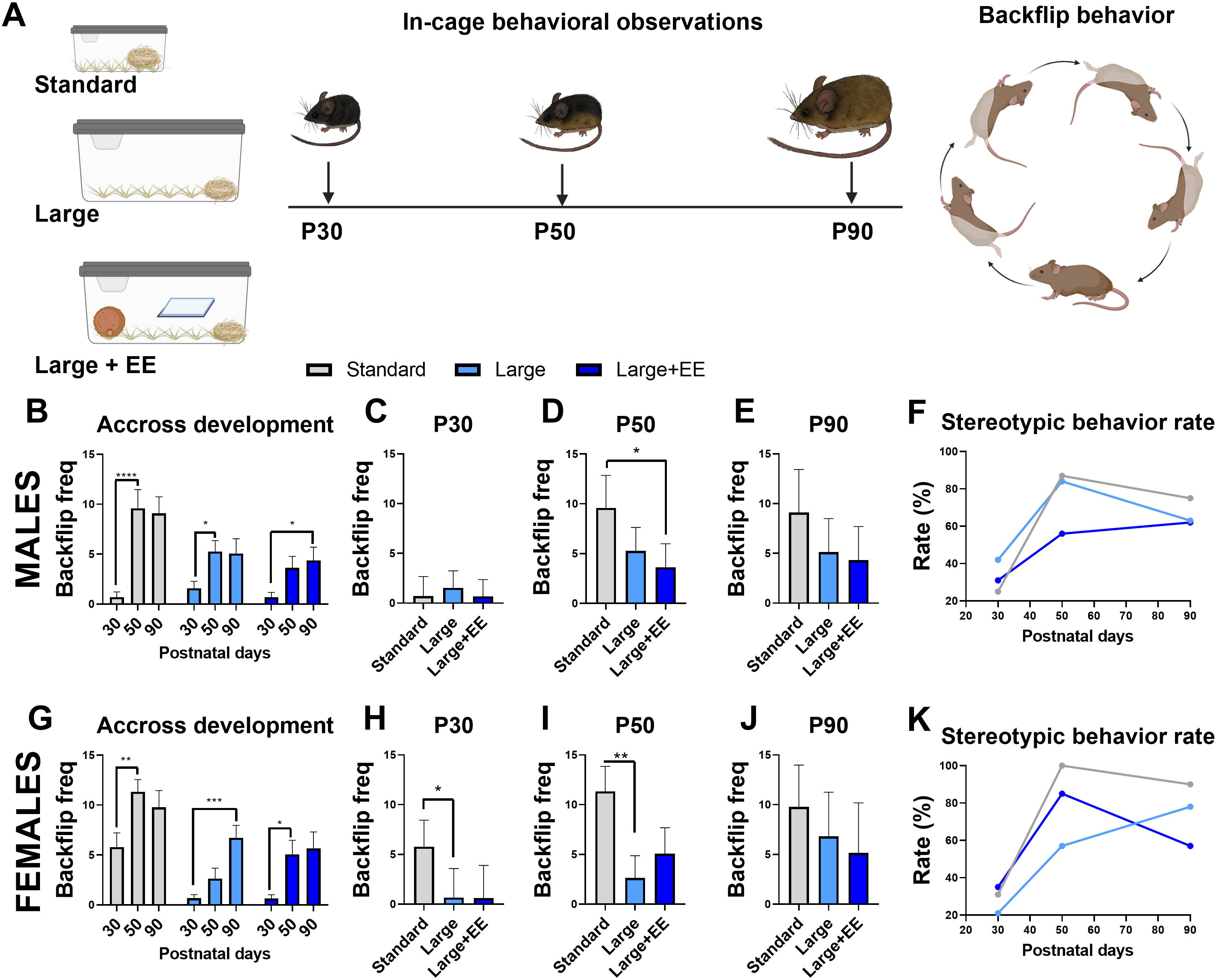
Backflipping behavior across development. Experimental timeline (A). Stereotypic behavior increases with age across experimental conditions in males (B) and females (C) (repeated 2-way ANOVAS). In males, large+EE reduces frequency of stereotypic behavior at P50, but not P90 (C-E) (nested 1-way ANOVA). In females, large, but nor large+EE, reduces stereotypic behavior at P30 and P50 (H-J) (nested 1-way ANOVA). Percent of animals showing stereotypic behavior within each housing group (F,K) (n=10-19 per group).

### Individual welfare status assessment

For a subset of the animals, individual welfare status was assessed every 7 days since weaning (P30) until one week before behavioral assessments started (P86) using a protocol adapted from Spangenber and Keeling (Elin MF Spangenberg and Linda J Keeling, 2016). Briefly, weight, body condition (scale 1-5, table 1), presence of injuries, coat condition, and response to handling (urination, defecation) were recorded (table 2).

### Social defeat stress

Once mice reached adulthood (3 months old), they were exposed to social defeat stress as previously described (Trainor et al., 2011). Briefly, the focal mouse was introduced into the home cage of a territorial same-sex conspecific for a duration of 7 minutes or until it was attacked 7 times (whichever occurred first). Immediately after each defeat episode was completed, the mice were returned to their home cages. Social defeat resulted in no physical wounds.

### Social Interaction test

Animals were tested in the social interaction test (Trainor et al., 2011) before (7 days prior) and after (7 days) exposure to social defeat stress (fig. 3A). Briefly, the social interaction test consisted of three 3 minute phases during which the behavior of the focal mouse was recorded and automatically scored using Any-maze (Stoelting): open field, acclimation, and social interaction. During the open field phase, the mouse was placed in the center of an empty testing arena (80 × 63 × 60cm) and was allowed to freely explore. During this phase, automatic scoring included total distance traveled to assess locomotor activity and time spent in the center of the arena (within 8cm of the sides and 14cm of the ends), which was used as a measure of anxiety-like behavior. During the acclimation phase, an empty wire mesh cage was placed against one of the walls of the arena. During the interaction phase, the wire mesh cage was replaced by another identical one containing a novel same-sex conspecific (target mouse). During both the acclimation and interaction phase, automatic scoring included time spent within 8cm of the mesh cage (interaction zone) and time spent in corners of the arena. Immediately after the interaction phase was complete, the animals were returned to their home cage.

**Fig. 3.**
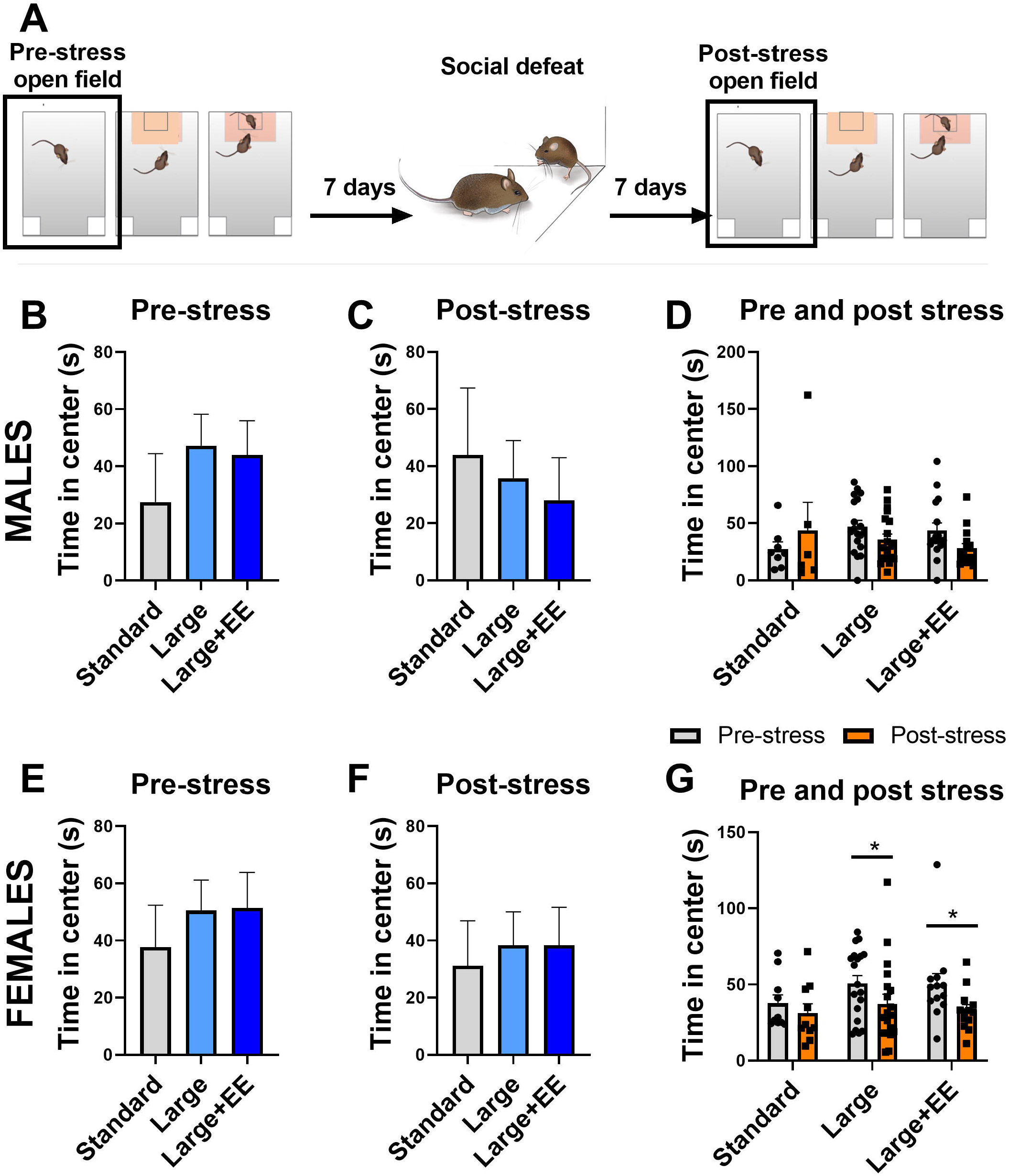
Behavior in the open field. A. Experimental timeline. Housing conditions do not affect time spent in the open field before or after stress in males (B,C) or females (E,F) (nested 1-way ANOVA). Stress does not affect time spent in the open field in males of any of the housing groups (D). Stress reduces time spent in the open field in females housed in large and large+EE, but not in standard cages (G) (repeated measures ANOVA).

### Statistical analyses

All analyses were done using Prism GraphPad®. Frequency of backflipping behavior was assessed using 1-way nested ANOVAs to compare housing conditions between groups at a given age (P30,P50,P90), and 2-way ANOVAsto assess effects of age on this behavior across groups. For the social interaction test, 1-way nested ANOVAs were used to compare time spent in the open field and social approach between housing conditions (pre and post stress were assessed separately), and 2-way ANOVAs to assess the effects of stress on this behavior across groups. For body weight and body condition, 2-way ANOVAs were used to assess this variable throughout development across groups. Finally, urination in response to handling was compared using nested 1-way ANOVA. Planned comparisons were performed if ANOVAs showed a significant effect, using Tukey for nested analyses and Sidak for 2-way ANOVAs to correct for multiple comparisons. Statistical significance was established at alfa<0.05 for all tests. An F-test was used to compare variances between groups.

## Results

### Backflipping behavior

In males, backflips were significantly affected by age in a housing-dependent fashion (Fig. 2B, repeated measures 2-WAY ANOVA main effect of age F2,90=20.51, p<0.0001, the main effect of treatment F2,45=4.23, p=0.02). Backflipping frequency increased from P30 to P50 in standard (d=2.1, p<0.0001) and large housing (d=0.6, p=0.03), but not in large+EE, which showed an increase from P30 only by P90 (d=0.6, p=0.03). When comparing backflipping behavior within age groups there was a trend for treatment to have an effect at P50 in males (Fig.2D, nested 1-way ANOVA F2,12=3.2, p=0.07). Post-hoc comparisons showed that larger+EE cages reduced backflip frequency compared to standard housing (d=0.96 p=0.03). No effects of housing were found in males at P30 or P90. Similar to males, in females, backflipping behavior was affected by age depending on housing conditions (Fig. 2F, mixed-effects model main effect of age F2,75=14.83, p<0.0001, the main effect of treatment F2,45=18.16, p<0.0001). Backflipping frequency increased from P30 to P50 in standard (d=1.21, p=0.009) and large+EE (d=1.56, p=0.03), but not in large cages, which showed an increase only by P90 (d=1.56, p=0.007). Comparisons within age groups revealed a main effect of housing conditions at P30 (Fig. 2G, nested 1-way ANOVA F2,30=3.9, p=0.03) and at P50 (Fig. 2H, nested 1-way ANOVA F2,30=5.4, p=0.01). Planned comparisons revealed that large cages, but not large+EE, significantly reduced backflipping behavior compared to standard at both P30 (d=1.13, p=0.03) and P50 (d=1.26, p=0.003).

### Open field

There were no significant differences between housing conditions in the time spent in the open field when comparing housing conditions before (Fig. 3 B,E) or after (Fig.3 C,F) stress in males or females (nested ANOVAs). When comparing the behavior before and after stress within groups (repeated measures 2-way ANOVAs), only in females there was a main effect of stress (Fig. 3G, F1,39=12.78, p<0.001). Planned comparisons showed that stress significantly reduced time spent in open field in females housed in large (d=0.5 p=0.03) and large+EE (d=0.52 p=0.02), but not standard cages.

### Acclimation

There were no effects of housing conditions or stress on time spent investigating an empty cage in males (fig.4B,C,D). Housing conditions did affect this behavior in unstressed females (Fig. 4E, nested one way ANOVA F2,40=5.478): standard cage females showed significantly less time exploring the cage compared to large (d=0.8, p=0.042) and large+EE (d=1.5, p=0.004) housed females. There were no differences between housing conditions in females after stress (fig. 4F). Interestingly, when comparing the effects of stress on time exploring an empty cage (fig. 4G, repeated measures 2-way ANOVA main effect of stress F1,42=5.75, p=0.02), we found that stress increased time exploring the object in standard (d=1.34, p=0.005), but not large or large+EE housed females.

**Fig 4.**
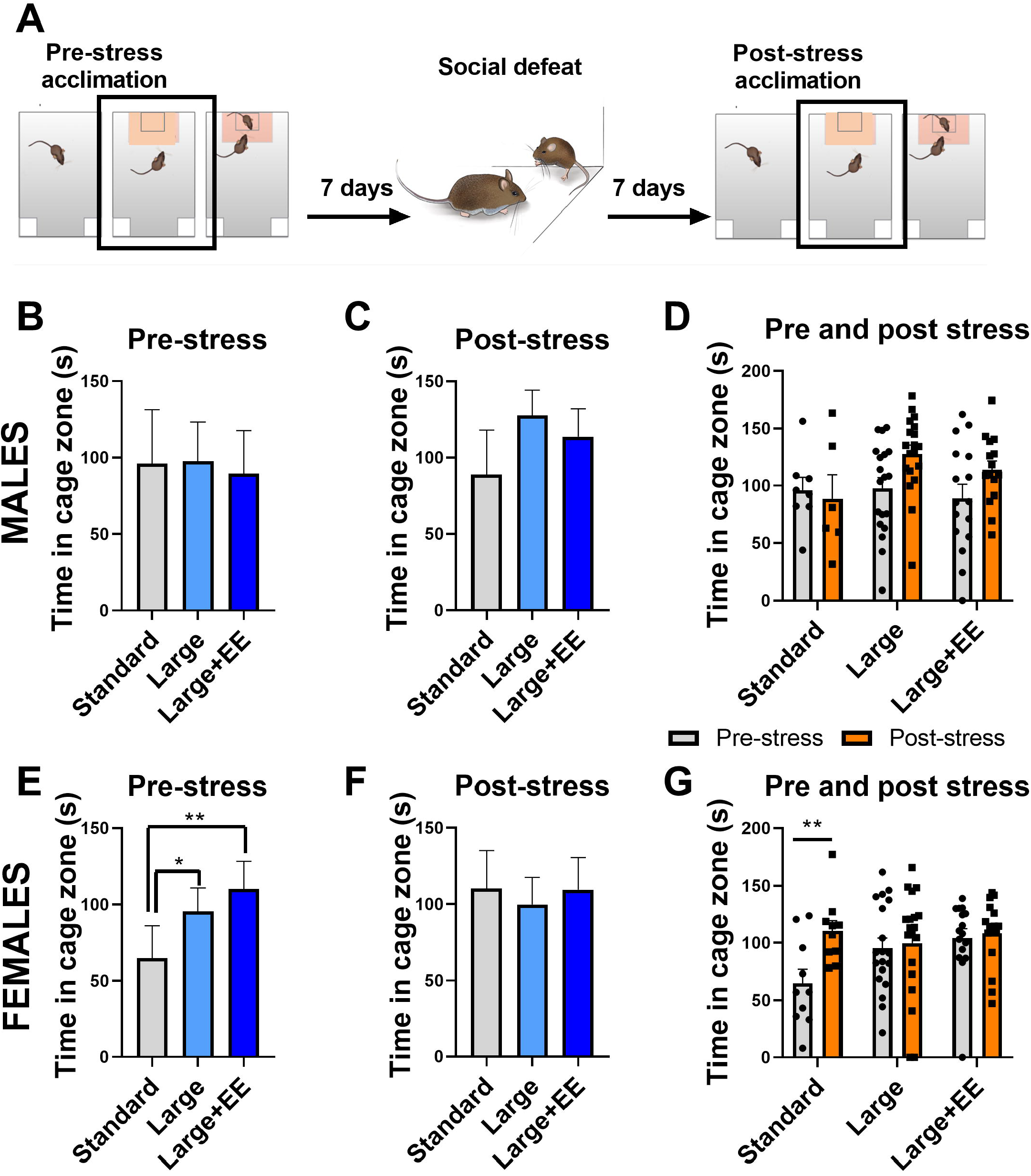
Behavior during acclimation phase (target absent) A. Experimental timeline. Housing conditions did not affect time spent in proximity to an empty cage in males either before (B) or after stress (C). Unstressed females housed in standard cages spent less time in proximity to an empty cage compared to females housed in large and large+EE (E), but housing had no effect on this measure after stress (F) (nested 1-way ANOVA). In males, stress had no effect on time spent in proximity to an empty cage in any of the housing groups (D). In females housed in standard, but not large or large+EE, stress increased time in cage during acclimation (G, repeated measures ANOVA).

### Social interaction behavior

In males, there were no effects of housing conditions on social approach behavior. All males showed high levels of social interaction both 7 days before and 7 days after stress (fig.4 B,C,D). On the contrary, pre-stress females showed different levels of social interaction depending on housing conditions (Fig. 5E, nested ANOVA main effect of treatment F2,12=4.05, p=0.04), with females in standard cages showing significantly less time in social approach than females in large+EE cages (d=1.1, p=0.02). Importantly, an F-test revealed that, compared to females housed in standard housing, females in large+EE conditions showed significantly less interindividual variance in the time spent in social approach (standard deviation 51.6 vs. 24.5, p=0.006). Thus, we repeated the analysis after a square root transformation to equalize variances, but the main conclusions did not change (main effect of treatment p=0.048, standard vs large+EE p=0.038). There were no differences between groups in females post-stress. Critically for our research program, social defeat stress reduced time spent in social approach in females of all groups (Fig. 5G, repeated-measures 2-way ANOVA main effect of stress F1,40=51.98, p<0.0001). This replicates previous findings showing that females, unlike males, are susceptible to showing long-term social deficits after exposure to social defeat stress (Duque-Wilckens et al., 2020, 2018; Steinman et al., 2016; Trainor et al., 2011), and shows that the effect is robust and persists regardless of housing conditions. Interestingly, the effect of stress on social approach was the largest in females housed in large+EE (d=2.2, p=0.0001), followed by large (d=1, p=0.0003), and standard housing (d=0.83, p=0.002).

**Fig. 5.**
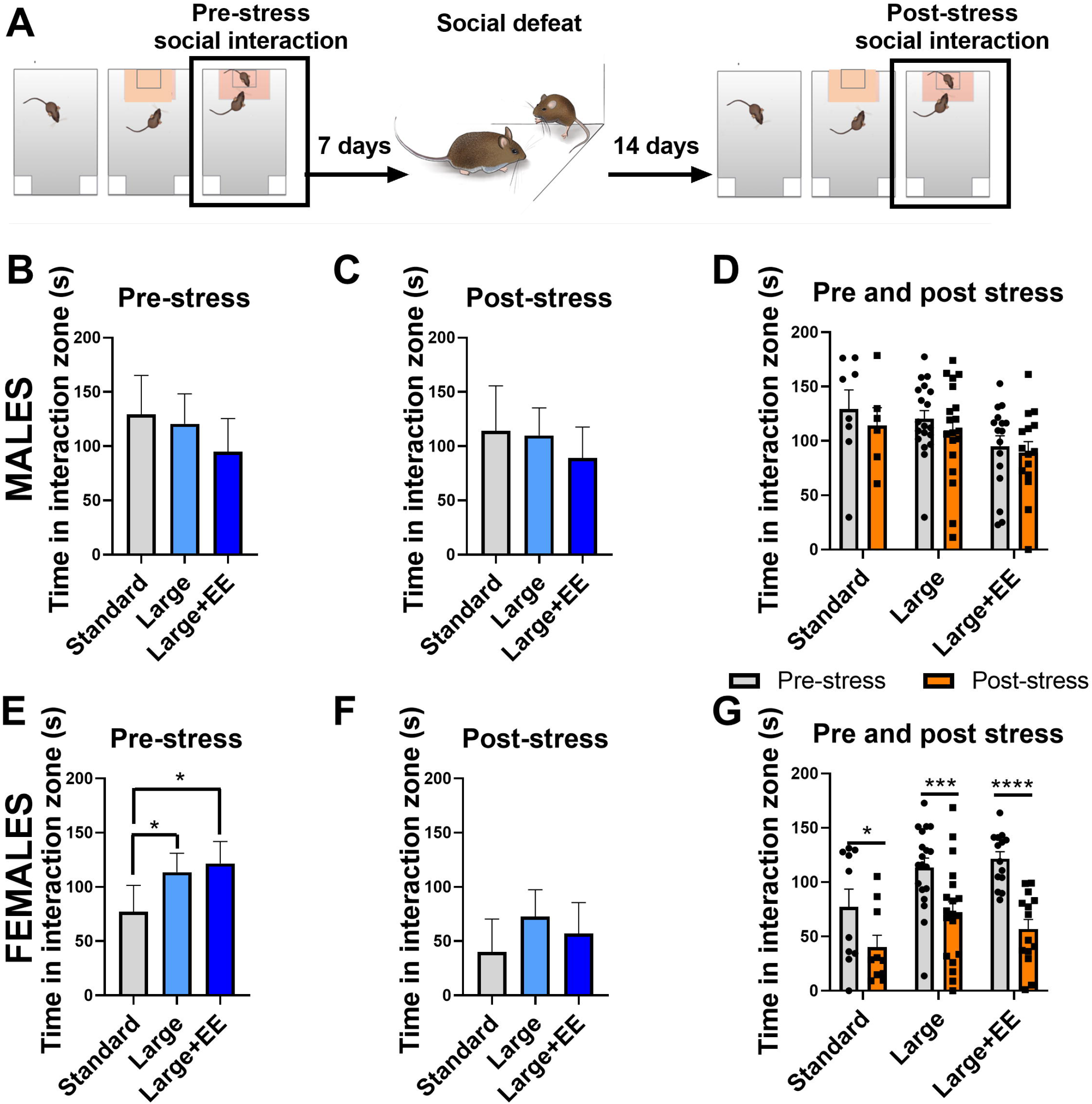
Behavior during interaction phase (target present) A. Experimental timeline. Housing conditions did not affect time spent in proximity to the social stimulus in males either before (B) or after stress (C). Unstressed females housed in standard cages spent less time in proximity to a social stimulus compared to females housed in large and large+EE (E), but housing had no effect on this measure after stress (F) (nested 1-way ANOVA). In males, stress had no effect on time spent in proximity to the social stimulus in any of the housing groups (D). In females of all housing groups, stress increased time in cage during social interaction, but the effect size was smallest in the standard housed females (G, repeated measures ANOVA).

### Use of enrichment

All animals except 3 males used the enrichment item at least once during the three 5 minutes in cage observations. There was a main effect of age on use of enrichment in males (1-way ANOVA main effect of age F2,47=5.8, p=0.005, fig. 5A) and females (1-way ANOVA main effect of age F2,32=6.0, p=0.006, fig. 5E), although in opposite directions: in males, the use of enrichment gradually increased with age, with use of enrichment at P30 being significantly lower that use of enrichment at P50 (d=1.0, p=0.03) and P90 (d=1.1, p=0.006). In females, on the contrary, the use of enrichment reduced from P30 to P90 (d=1.5, p=0.004).

### Use of Enrichment

In males, the use of enrichment items increased throughout development (Fig. 6A, 1-way ANOVA F2,47=5.8, p=0.005). Compared to P30, males used enrichment more frequently both at P50 (p=0.03) and P90 (p=0.006). Interestingly, females showed the opposite: frequency of enrichment use decreased with time (Fig. 6B, 1-way ANOVA F2,32=6, p=0.006), compared to P30, females used enrichment less frequently at P90 (p=0.004).

**Fig. 6.**
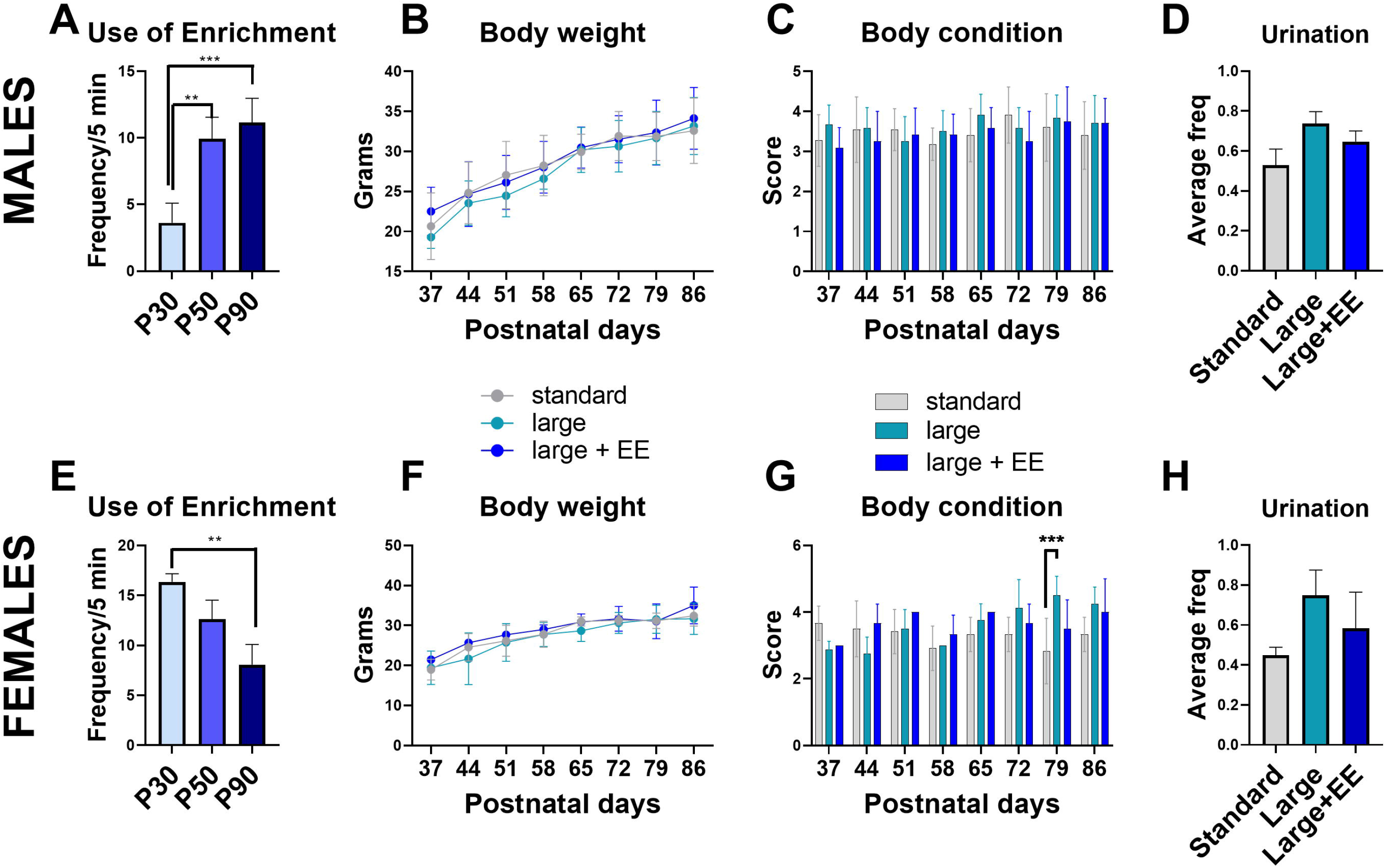
Use of enrichment increased with age in males (A), but decreased in females (E) (1-way ANOVA). There were no differences in body weight between housing groups in males (B) or females (F). There were no differences in body condition between housing groups in males (C). In females large increased body condition score compare to standard at PN79 (2-way repeated measures ANOVA). There were no effects of housing on urination in response to handling in males (D) or females (H).

### Welfare and body condition analyses

We did not see the effects of housing conditions on body weight (Fig. 6B,F) or urination frequency (Fig.6D,H) in response to handling in males or females. In females, housing affected body condition but only in older mice (Fig. 6G, 2-way ANOVA F14, 70=2.83, age x housing interaction p=0.002). Body condition scores were higher in females housed in large cages compared to standard caging at PN 49 (p=0.0002) and marginally higher at PN 56 (p=0.06). In males, housing conditions had no significant effect on body weight or body condition across development (all p’s > 0.26). We also did not detect any presence of injuries or differences in coat condition. Animals from all groups showed a similar frequency of urination in response to handling (Fig. 6 D,H). No defecation in response to handling was recorded.

## Discussion

The present studies show that increasing housing space as well as social and environmental complexity in California mice can delay the development of stereotypic behavior. Critically, this reduction in stereotypy is accompanied by increased effect sizes of stress in an established model for social anxiety. These results suggest that increased cage size and enrichment enhances welfare for mice while simultaneously enhancing the quality of behavioral experiments.

### Housing conditions on expression of stereotypic behavior

We found that the effects of housing on stereotypic behavior are age and sex-dependent. In males of all housing conditions, backflipping frequency was very low at P30 and significantly increased at P50 in standard and large housing groups, but not in large+EE. In this group, the backflipping behavior did increase, but only by P90. A similar effect has been reported in deer mice, where increased space and environmental complexity delayed the appearance of stereotypic behavior compared to standard housed animals (Powell et al., 1999). This suggests that, although increased space and social/environmental complexity had a positive impact on stereotypic behavior during earlier stages of development, it was not sufficient to inhibit its appearance later in life. The results in California mice could be partly explained by the fact that this species initiates dispersal from their parents’ nests betweenP75-85 in the wild (Ribble, 1992). Thus, between P50 and P90, a motivational shift could render the available space and environmental complexity in large+EE insufficient for the individuals to perform age-specific behaviors essential to survival and reproduction in the wild, resulting in increased backflipping (Garner, 2005).

A similar housing-dependent delay of stereotypic behavior was seen in females. Nonetheless, in this case large, but not large+EE, shifted the increase from P50 to P90 compared to standard housing. This suggests that, while increased space and social complexity is effective at delaying stereotypies in females, the addition of environmental enrichment somehow negates this effect. It is possible that added enrichment items increased intra-cage competition, although this was not measured in the present study. Some studies in C57 mice have previously reported that the addition of environmental resources can result in increased aggression (Barnard et al., 1996; Haemisch and Gärtner, 1997). If this is the case, it would be intriguing to understand why competition develops as a response to environmental enrichment only in females, as both female and male California mice show high levels of territorial aggression towards same-sex conspecifics (Rieger et al., 2019; Trainor et al., 2011).

Another important observation in the current studies is that females housed in standard housing conditions showed higher frequency of stereotypic behavior already at P30, in contrast to males who only showed an increase in this behavior by P50. This could be associated with differences in sexual maturity: female California mice molt their juvenile coat earlier than males (Wright et al, unpublished). Future studies assessing the underlying neurobiology of stereotypic backflipping behavior could shed more light on this question.

### Effects of housing conditions on anxiety-like and exploratory behaviors

In females both large and large+EE affected behavior in a context-dependent manner, in contrast, no effects of housing or stress were observed in males. Under baseline (non-stress) conditions, large and large+EE increased the time females spent in non-social and social exploration without affecting time spent in the open field. This suggests that elevated motivation, rather than reduced anxiety, underlies the effects of housing on these behaviors. Previous studies have found that environmental enrichment has profound effects on the physiology of ventral striatum, a key brain area underlying motivation and reward processing. For example, microdialysis studies in rats showed that environmental enrichment results in elevated levels of extracellular striatal dopamine (Segovia et al., 2010), and studies in mice have shown that enrichment alters the expression of genes key to striatal signaling, including upregulation of Brain-derived neurotrophic factor (BDNF) (Bezard et al., 2003), and the transcription factor delta-Fos B (Solinas et al., 2009).

The fact that both large and large+EE had similar effects on exploratory behaviors suggests that increased space and social complexity, in the absence of environmental enrichment, are sufficient to trigger an effect on motivated responses. Studies in female C57 mice showed similar results: just the social component of enriched environments could explain an increase in both non-social and social exploration time, which was correlated with an increased number of doublecortin expressing cells in the hippocampus, indicating increased neurogenesis (Moreno-Jiménez et al., 2019).

We were particularly interested in learning whether housing conditions would interfere with the sex-specific effects of defeat stress on social interaction behavior that we consistently see in our research program (Duque-Wilckens et al., 2018; Greenberg et al., 2014; Trainor et al., 2011). Remarkably, we found that social defeat reduces time spent in social approach in females of all housing groups without affecting this behavior in males. These results show that female-biased susceptibility to social defeat stress is a very robust phenotype in California mice, which provides even stronger support for using this model as a means of understanding female-biased psychiatric disease. Further, we found that both large and large+EE increase the magnitude of the effects of stress on social approach by reducing the within-group variance in females. This is very relevant as it could translate into fewer animals needed to achieve statistical significance, favoring the second of the three Rs principles underpinning the humane use of animals in scientific research (van Luijk et al., 2013). Overall, the current studies show that larger cages and increased social and environmental complexity effectively delay the appearance of stereotypic behavior. Importantly, these reductions in stereotypic behaviors are associated with reduced variability and increased effects size in behavioral experiments focusing on social anxiety-related behaviors. These results highlight the benefits of paying close attention to housing conditions to improve animal welfare and quality of science.

## Acknowledgments

The authors acknowledge Cindy Clayton for veterinary care, and Amanda Kentner for advice on experimental design. This work was supported by Grants for Laboratory Animal Science (GLAS) program of the American Association for Laboratory Animal Science (AALAS) to NDW, NIH R01MH121829 and NSF IOS 1937335 to BCT

## Notes

### Competing Interest Statement

The authors have declared no competing interest.

