## Supplementary material for "Environmental enrichment delays the development of stereotypic behavior and reduces variability in behavioral experiments using California mice (*Peromyscus californicus*)": Table 1

Table 1. Body condition score

| **Body condition score** | **Description** |
| --- | --- |
| 1 | Spine is prominent, animal is emaciated |
| 2 | Spine somewhat prominent, animal is underweight |
| 3 | Spine not prominent, animal is at a healthy weight |
| 4 | Spine only palpable with pressure, animal is overweight |
| 5 | Spine is obscured by fat and flesh, animal is obese |

Table 2. Behavior in response to handling

| **Measure** | **Description** |
| --- | --- |
| Urination | Excretion of urine during handling yes/no |
| Defecation | Excretion of feces during handling yes/no |
| Injuries | Presence/absence |
| Coat condition | Good/poor |
